# Unconventional Small Molecule MNP-021 Protects Neuronal and Glial Function from Diabetes-Associated Glucotoxicity and Neuroinflammation

**DOI:** 10.64898/2026.01.29.702298

**Authors:** Alessandro De Carli, Carolina Filipponi, Beatrice Polini, Veronica Sancho-Bornez, Emy Basso, Fabio Filippini, Angela Dardano, Francesca Sardelli, Simona Daniele, Martina Contestabile, Giuseppina Emanuela Grieco, Francesco Dotta, Guido Sebastiani, Maria Grazia Chiellini, Mauro Pineschi, Michele Lai, Giuseppe Daniele

**Author notes:** **Corresponding Author:** Giuseppe Daniele, MD, PhD, Department of Clinical and Experimental Medicine, University of Pisa, Italy. Authors contributed equally (Alessandro De Carli, Carolina Filipponi).

## Abstract

**Background:** Diabetes-associated neurodegeneration is amplified by methylglyoxal (MGO)-driven dicarbonyl stress linking hyperglycemia to neuronal insulin resistance and maladaptive neuroinflammation. We tested the neuroprotective activity of MNP-021, a non-electrophilic TRPA1 modulator, in neurons and glial cells *in vitro*.

**Methods:** SH-SY5Y neurons were pretreated with MNP-021 and challenged with MGO, then profiled by high-content imaging, RNA-seq, Seahorse OCR/ECAR, glycolytic stress assays and AKT/ERK/CREB immunoblotting ± insulin. In parallel, HMC3 glial cells were treated with MNP-021, exposed to LPS/TNF-α or Aβ_(25–35)_ and tested for viability and inflammatory markers by ELISA and qRT-PCR.

**Results:** MGO increased nucleus-to-cytoplasm area ratio by 49% and dysregulated glucose handling, increasing 2-NBDG uptake by ∼25%, with GLUT1/GLUT4 membrane redistribution; MNP-021 normalized morphology, uptake, and transporter localization without cytotoxicity up to 10 µM. RNA-seq identified 754 MGO-deregulated genes, including ISR/metabolic nodes (GCK, SESN2, PHGDH/PSAT1, PCK2); MNP-021 buffered stress-induced transcription with limited baseline effects, remodeled mitochondrial redox readouts consistent with controlled ROS signaling, while improving mitochondrial content/architecture and blunting stress-evoked compensatory glycolysis. MNP-021 restored pro-survival signaling (pAKT/pERK and nuclear pCREB), including insulin responsiveness during MGO exposure. MNP-021 reduced IL-6/TNF-α release while increasing IL-10 and ARG1 (∼1.9-fold vs LPS/TNF-α) in HMC3 glial cells, shifting them toward a pro-resolving IL-10/ARG1 program with reduced Aβ_(25–35)-_evoked cytokine release with GLP-1 remaining very low (≤10 pg/mL) and not significantly increased in this system.

**Conclusions:** MNP-021 coordinates transcriptomic restraint, transporter-level glucose handling, mitochondrial resilience, and pro-survival/pro-resolving signaling across neuron–microglia compartments, supporting TRPA1-tuned small-molecule modulation as a candidate strategy against dicarbonyl-linked neuro-metabolic stress.

## 1. Introduction

Diabetes mellitus is a chronic metabolic disease defined by persistent hyperglycemia, with clinically relevant involvement of peripheral and central nervous systems. Long-term epidemiological evidence links diabetes to accelerated cognitive decline over decades, consistent with dysmetabolism-driven brain vulnerability accumulating across the life course^1^. Despite major advances in glucose- and weight-lowering therapies, evidence that current metabolic treatments consistently prevent or slow cognitive deterioration remains limited^2,3^. Together, these observations suggest that improving metabolic endpoints alone may not sufficiently address the cellular mechanisms coupling hyperglycemia to neurodegenerative trajectories. Mechanistically, diabetes-related neurodegeneration reflects convergence of oxidative and dicarbonyl stress, mitochondrial dysfunction, inflammation, and advanced glycation end-product (AGE) accumulation, culminating in neuronal dysfunction/apoptosis and glial reactivity^4,5^. Among reactive mediators elevated in hyperglycemia, methylglyoxal (MGO), a highly reactive glycolytic byproduct, accumulates in diabetes and is a key effector of glucotoxic dicarbonyl stress^5^ and altered Ca² /stress homeostasis^6^. MGO is also an endogenous activator of transient receptor potential ankyrin 1 (TRPA1), shown to drive nociceptor sensitization and pain, thereby connecting metabolic byproducts to excitability-linked pathology relevant to metabolic neuropathies^7^. Beyond excitability, MGO compromises insulin/trophic signaling independently of intracellular ROS formation, supporting “dicarbonyl-driven insulin resistance” at the IRS–PI3K interface with downstream consequences for AKT/ERK-dependent survival^8^. Given the broad role of ERK1/2 pathways in diabetes-related pathogenesis and therapeutic relevance, disruption of AKT/ERK signaling is particularly consequential for neuronal resilience under metabolic stress^8^. Omics data further indicate that MGO acts as a systems-level disruptor, driving remodeling of metabolic networks, stress responses, cytoskeletal organization, and mitochondrial function alongside adaptive redox programs^9^.

Glial biology provides an essential second compartment in diabetes-associated neurodegeneration. Microglia can amplify neuronal dysfunction through inflammatory cytokines, innate immune activation and chronic engagement is implicated across neurodegenerative diseases^10,11^. Microglial states are stimulus- and context-dependent, spanning detrimental pro-inflammatory programs and pro-resolving phenotypes that support repair, making microglia a rational target for qualitative modulation rather than blanket suppression^12–14^.

In Alzheimer’s disease, amyloid-β (Aβ) intersects with immune metabolism and microglial function, and metabolic reprogramming failure has been linked to microglial dysfunction^14^. Aβ-driven inflammatory toxicity can be modeled and pharmacologically modulated in human HMC3 glial cells, providing a tractable system to probe microglial responses under inflammatory/amyloid challenge^15^. Collectively, these observations support a feed-forward loop in which hyperglycemia-driven carbonyl stress potentiates mitochondrial dysfunction and immune activation, and glial inflammation further destabilizes neuronal metabolic homeostasis^4,5,10,11,14,15^.

Within this framework, TRPA1 has emerged as a molecular node at the interface of oxidative/carbonyl stress, inflammation, and metabolism^16^. Pineschi and colleagues described non-electrophilic TRPA1 modulators that stimulate Glucagon-like Peptide-1 (GLP-1) release in enteroendocrine cells and protect human neurons from MGO- or high-glucose–induced apoptosis, suggesting that tuned TRPA1 engagement can yield beneficial outcomes distinct from classical electrophilic TRPA1 agonism^17^. These findings motivated investigation of lead compound 4b, here renamed MNP-021, a 1,3-diaza-4-oxabicyclic[3.3.1]nonene derivative, as a candidate scaffold with metabolic modulatory and neuroprotective potential^17^. The aim of this study was therefore to define the protective profile of MNP-021 in human neuronal (SH-SY5Y) and microglial (HMC3) models exposed to glucotoxic and inflammatory stress, using a multi-platform strategy linking phenotype to mechanism. By addressing both neuronal and glial compartments, this work seeks to position MNP-021 as a candidate therapeutic approach for diabetes-associated neurodegeneration and to provide a rationale for subsequent *in vivo* and translational development.

## 2. Methods

### 2.1 SH-SY5Y cells and treatments

SH-SY5Y neuroblastoma cell line was cultured in DMEM/F12 (Gibco, Thermo Fischer Scientific, USA) supplemented with 2 mM L-glutamine, non-essential amino acids, and 10% fetal bovine serum (FBS, Sigma) at 37°C and 5% CO2. Cells were tested for mycoplasma contamination.

Experiments were conducted exclusively on differentiated mature-like neurons, following a 6-days protocol in medium containing 10 µM retinoic acid (RA). On day six, all experiments were carried out according to the treatment protocol, which included four experimental groups: cells treated with 400 µM methylglyoxal (MGO) (Sigma, M0252) for 4 hours, as stress positive control; cells treated with 10 µM MNP-021 for 48 hours; cells treated with a combination of both: for the last 4 of 48 hours, cells received 400 µM methylglyoxal + 10 µM MNP-021; and a control group treated with DMSO for 48 hours.

### 2.2 RNA extraction and Transcriptomics libraries preparation and QC on SH-SY5Y

Total RNA was extracted through DirectZol RNA microPrep Kit (Zymo Research #R2062) following the manufacturer’s instructions.

RNA QC was performed by QUBIT 3.0 fluorometer with RNA HS assay (Q33223), and Tape Station 4150 (Agilent Technologies, RNA ScreenTape #5067-5576 and reagents #5067-5577 #5067-5578) to assess concentration and RNA integrity (ensuring RIN value ≥7.0), respectively.

RNA-seq Libraries were prepared using 100 ng of input RNA through NEBNext rRNA Depletion Kit v2 (Human/Mouse/Rat) with Beads (#E7405), NEBNext® Ultra II Directional RNA Library Prep Kit for Illumina® with Sample Purification Beads (#E7765), and NEBNext Multiplex Oligos for Illumina (Unique Dual Index UMI Adaptors RNA Set 1 #E7416) following manual’s instructions.

Library QC and concentration were evaluated through QUBIT 3.0 fluorometer with DNA HS 1X assay (Q33230), and TapeStation D1000 ScreenTape (5067-5582 and reagents kit 5067-5583).

Libraries were normalized to 3nM, pooled, and sequenced on Novaseq6000 platform at a final concentration of 600 pM.

### 2.3 Bioinformatic analysis

Paired-end FASTQ files were trimmed, removing adapters and low-quality reads. Trimmed FASTQ files were then aligned using the STAR algorithm (Version 2.7.11b) with the Genome Reference Consortium Human Build 38 patch release 14 (GRCh38.p14) as reference. The resulting BAM files were indexed with Samtools (Version 1.22.1) and PCR duplicates were removed using the UMI-tools algorithm (Version 1.1.6).

Transcript quantification was performed with the featureCounts algorithm (Version 2.1.1), utilizing the GRCh38.p14 GTF annotation file as a reference, producing a raw counts matrix.

The DESeq2 (Version 1.49.4) package’s variance stabilizing transformation (VST) was applied to the raw counts matrix to achieve homoscedastic data for a Principal Components Analysis (PCA) to evaluate samples segregation based on experimental condition (CTR, MNP-021, MGO or MNP-021+MGO) within each experimental setup.

Low-expression genes were filtered out from raw count matrix for all samples, retaining transcripts with at least one Counts Per Million (CPM) in at least one sample, for further analyses using the EdgeR package (Version 4.6.3). DESeq2 was used to normalize the filtered count matrices by library size and RNA composition, and to identify differentially expressed (DE) genes across experimental conditions, using the median of ratios method and Wald’s test (paired), respectively. The resulting P-values were adjusted using the Benjamini-Hochberg method to correct for multiple comparisons. Transcripts, with adjusted P-values (padj) < 0.05 and log2FC<-0.6 (downregulated) or log2FC>0.6 (upregulated), were classified as differentially expressed between conditions.

DE genes were compared between: MNP-021 vs CTR, MGO vs CTR, MNP-021+MGO vs MGO. Resulting DE genes were subjected to pathways enrichment analysis using DAVID (version 6.8) algorithm. Biological processes (BP) identified were considered statistically significant with p-value corrected for False Discovery rate (FDR) <0.05.

### 2.4 Immunofluorescence on SH-SY5Y for cell morphology and GLUT transporters localization

Cells were fixed using 4% paraformaldehyde (PFA). Then, cells were permeabilized with PBS Ca^2+^Mg^2+^ Triton X-100 0.5%. Subsequently, cells were blocked with PBS Ca^2+^Mg^2+^ Triton X-100 0.3 % and FBS 3% and stained for GLUT1 (Thermo Fisher Scientific, MA5-31960), 1:500, and GLUT4 (Thermo Fisher Scientific, MA5-17176), 1:400.

To label primary antibodies, we used goat anti-mouse Alexa Fluor 568 (Invitrogen, Cat# A11004, 1:1000) and goat anti-rabbit Alexa Fluor 647 (Invitrogen, Cat# A21245, 1:1000) for GLUT4 and GLUT1, respectively. DAPI (Sigma Aldrich, St. Louis, MO, USA, 1:1000) and phalloidin 488 (Abcam, Cat# ab176759, 1:2000) were used to stain nuclei and cytoplasm. respectively. Images were acquired by Operetta CLS platform (Revvity, Waltham, MA, USA) using 40x magnification.

### 2.5 Glucose uptake analysis with 2-NBDG

After the treatments described above, we administered 0.2 µg/µL 2-NBDG (Invitrogen, Thermo Fisher Scientific, N13195), a fluorescently labeled deoxyglucose analog, dissolved in low-glucose Opti-MEM medium (Gibco, Thermo Fisher Scientific, USA) for 30 minutes at 37°C. Hoechst 33342 (Thermo Fischer Scientific, Cat# H1399, 1:1000) and CellMask Deep Red (Invitrogen, Thermo Fischer Scientific, Cat# C10046, 1:1000) were used to label nuclei and cell membrane. Images were acquired by Operetta CLS platform (Revvity, Waltham, MA, USA) set in normal growing conditions, using 40x magnification.

### 2.6 Mitochondrial analysis

After treatments described above, the cells were stained with 1 µM of MitoSOX Red superoxide indicator (Thermo Fischer Scientific, Cat# M36009), following the manufacturer’s protocol, to label Superoxide (SOX) in mitochondria. The staining was performed on living cells, and nuclei and cell membranes were labelled as above. Moreover, mitochondria were stained using 100 nM of MitoTracker Green FM (Thermo Fischer Scientific, Cat# M7514) for 30 minutes to assess the morphology of mitochondria. After staining, images were acquired by Operetta CLS platform (Revvity, Waltham, MA, USA) set in normal growing conditions, using 63× magnification.

### 2.7 Analyses of High-content confocal images

Images were processed using Harmony 5.1 Software (Revvity, Waltham, MA, USA). Each replicate was the mean of ≥ 1000 cells analyzed. The building blocks are available in the Supplementary Material section.

### 2.8 Oxygen consumption rate and extracellular acidification rate

Live oxygen consumption rate (OCR) and extracellular acidification rate (ECAR) were measured simultaneously using the Seahorse XFe24 Analyzer (Agilent).

SH-SY5Y cells were seeded in Seahorse XF well plates, coated with 10 μg/mL poly-D-lysine and 2 μg/mL laminin, using a complete DMEM/F12 medium.

After the differentiation protocol and treatments described above, the Seahorse measurements were performed following the manufacturer’s instructions. Optimal concentrations of oligomycin, FCCP, rotenone, and antimycin were determined through prior titration experiments.

### 2.9 Glycolysis stress test

In addition to the treatments described above, glucose oxidase (GOX) and untreated cells were included as a positive control for glycolytic activation and a baseline reference condition, respectively. ECAR was measured using the glycolysis stress test kit (Abcam, Cat# ab222946), performing sequential additions of free respiration buffer (FRB, baseline), oligomycin + glucose (glycolytic stimulation), and 2-deoxyglucose (2-DG) + glucose to inhibit glycolysis. Fluorescence readings were converted to acidification rates (H /h). ECAR was derived as: glycolytic capacity = Oligo − FRB, and non-glycolytic acidification = DG − Oligo.

### 2.10 Separation of the cell components: cytosol, membrane, and nucleus

Refer to the Supplementary Material section.

### 2.11 Protein expression analysis by Western Blot

Western blot analysis was performed on nuclear, cytoplasmic, and membrane fractions to evaluate the immediate effects of the compounds on the phosphorylation of ERK1/2 and AKT, their activated form, and the nuclear translocation of phosphorylated ERK1/2 (pERK1/2) and phosphorylated CREB (pCREB). We compared the effect of the compounds without and with the administration of 50 nM of insulin. Insulin was added for 30 minutes at the end of the treatments described above. Please refer to the Supplementary Material section for lysis and western blot procedures, and Supplementary Table 1 for antibody details.

### 2.12 Kinetic analysis by immunoenzymatic assay

A kinetic analysis was performed using an immunoenzymatic assay to quantify the levels of pAKT, pERK1/2, and pCREB, as well as total AKT and ERK1/2 (antibody conditions are described in Supplementary Table 2) after the treatments. 5 × 10□cells were treated as previously described with varying exposure times (180, 120, 60, and 30 minutes). Then, cells were fixed with 2% formaldehyde. Subsequently, a quenching solution (10% hydrogen peroxide and 100× sodium azide in 1× PBS) was added and incubated for 20 minutes. Then, cells were blocked in PBS1% BSA and 0.1% Triton X-100, and were incubated in primary antibody (described in Supplementary Table 2) diluted 1:300 in 1% BSA and 0.03% Triton X-100. Subsequently, cells were incubated with the secondary antibody for 2 hours. Finally, after 3,3′,5,5′-Tetrametil-benzidina substrate (TMB) addition and color development, absorbance was measured at 450 nm. Subsequently, we normalized the cell density as described in the Supplementary Material section.

### 2.13 HMC3 cells and treatments

The human microglial clone 3 cell line (HMC3) (ATCC® CRL-3304™) was cultured in high-glucose EMEM supplemented with 10% fetal bovine serum (FBS), penicillin (100 U/mL), and streptomycin (100 μg/mL) (Sigma-Aldrich, Milan, Italy), and maintained at 37 °C in a humidified atmosphere containing 5% CO□. Lipopolysaccharide (LPS; Escherichia coli O111:B4), tumor necrosis factor-α (TNF-α), and amyloid β-peptide _25–35_ (Aβ_25–35_) were purchased from Sigma-Aldrich (Milan, Italy). Vehicle-treated cells were used as controls in all experiments.

### 2.14 Cell viability assay (MTT) on HMC3 cells

Please refer to the Supplementary Materials section.

### 2.15 Release of inflammatory mediators from stimulated HMC3 cells

HMC3 cells were exposed to different pro-inflammatory stimuli. For LPS/TNFα-induced inflammation, cells were pre-treated with MNP-021 for 1 h and then stimulated with LPS (10 μg/mL) and TNF-α (50 ng/mL) for 24 h. For amyloid-induced inflammation, HMC3 cells were exposed to Aβ_25–35_ (10 μM) for 24 h; where indicated, cells were pre-treated with MNP-021 for 24 h before Aβ_25–35_stimulation. Vehicle-treated cells served as controls in all experimental conditions.

The concentrations of interleukin-6 (IL-6), interleukin-10 (IL-10), and tumor necrosis factor-α (TNF-α) were quantified using specific ELISA kits (IL-6, RAB0306; IL-10, RAB0244; TNF-α, RAB1089; Sigma-Aldrich, Milan, Italy), according to the manufacturer’s instructions.

### 2.16 RNA extraction and RT-qPCR analysis on HMC3 cells

Total RNA was extracted using the RNeasy Mini Kit (Qiagen, Cat# 74104, Hilden, Germany), according to the manufacturer’s instructions. RNA concentration was determined using a Qubit 3.0 fluorometer (Thermo Fisher Scientific, Wilmington, DE, USA). Then, 1 µg of total RNA was subsequently reverse transcribed using the iScript™ gDNA Clear cDNA Synthesis Kit (Bio-Rad, Milan, Italy). Gene expression analysis of MHC II, MCP1, and ARG-1 was performed by quantitative real-time PCR (RT-qPCR) using SYBR Green chemistry and the CFX Connect™ Real-Time PCR Detection System (Bio-Rad, Milan, Italy) with the following thermal profile: 95 °C for 30 s, followed by 40 cycles at 95 °C for 5 s and at 60 °C for 15 s. A melting curve analysis was included.

GAPDH was used as the endogenous reference gene, and relative mRNA expressions were calculated using the 2□^ΔΔCt^ method. Primer sequences were reported in Supplementary Table 3.

### 2.17 GLP-1 quantification by ELISA

Please refer to the Supplementary Materials section

### 2.18 Statistical analysis

Statistical analyses were performed using GraphPad Prism version 9.0 for Mac (GraphPad Software, San Diego, CA, USA), and significant differences among different treatments were calculated using ordinary One-way ANOVA followed by Dunnett’s or Tukey’s post hoc. All data are reported as mean ± SEM. Differences at p < 0.05 were considered significant.

## 3. Results

### 3.1 MNP-021 counteracts MGO neurotoxic effect without cytotoxicity in SH-SY5Y

Figure 1a outlines the experimental workflow used to assess the neuroprotective potential of MNP-021 in a diabetes-mimicking neuronal injury model, employing high-content confocal microscopy with automated image analysis. First, we characterized MNP-021 toxicity in SH-SY5Y cells using a dose-escalation design. As shown in Figures 1b–c, 10 µM represented the highest non-toxic concentration, consistent with Pineschi et al.^17^. SH-SY5Y-derived neurons were then pretreated with MNP-021 for 48 h and challenged with MGO (400 µM) for 4 h, after which cells were analyzed by high-content screening. Representative images and quantification of the morphological readout are shown in Fig. 1d–e. MGO induced a 49% increase in the nuclei-to-cytoplasm area ratio versus vehicle-treated cells. In contrast, MNP-021 treatment in the presence of MGO restored the nuclei/cytoplasm area ratio to values comparable to vehicle (Fig. 1d–e), supporting a neuroprotective effect of MNP-021.

**Figure 1.**
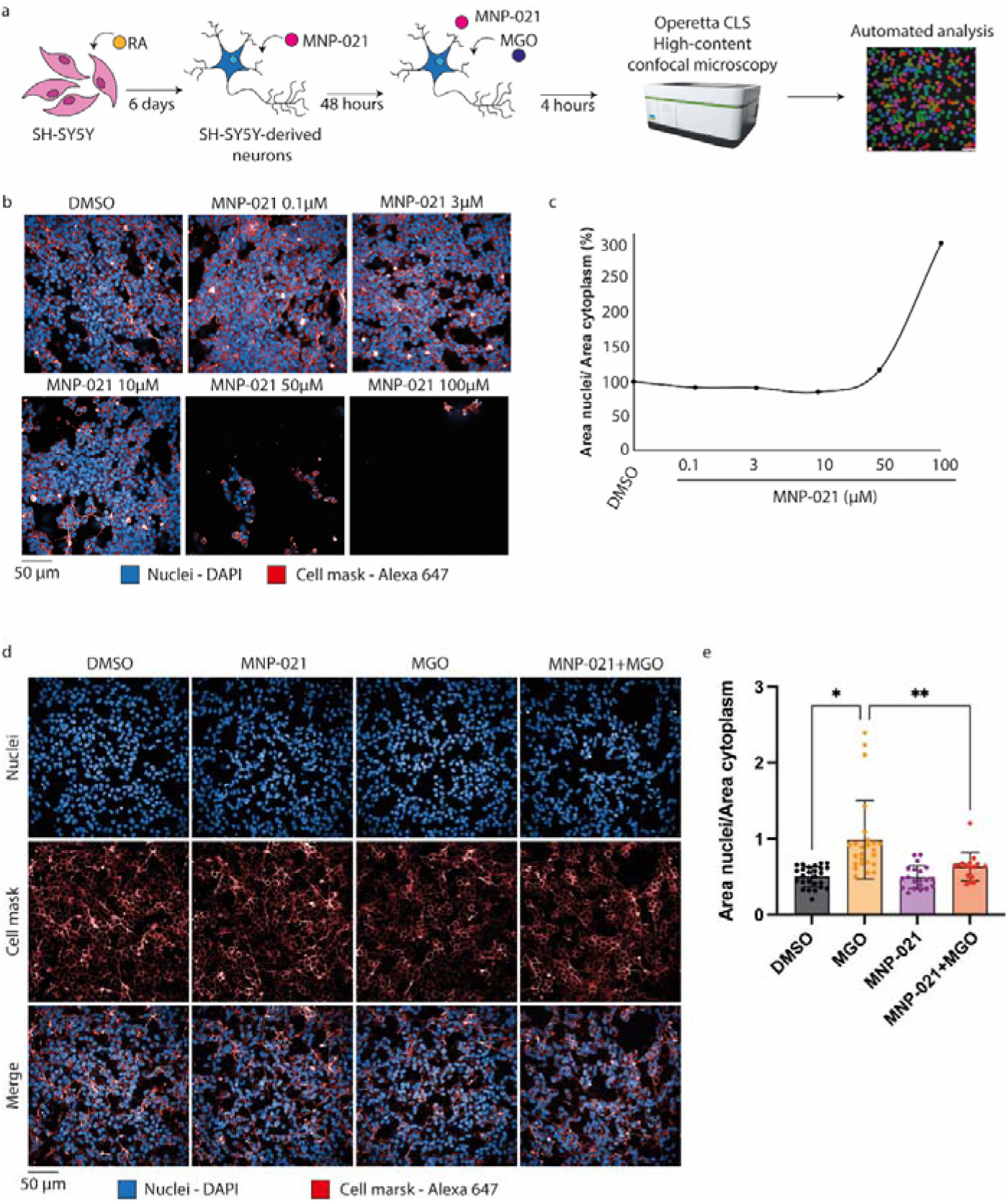
MNP-021 toxicity and neuroprotective effect on SH-SY5Y cells. a. Schematic representation of the experimental workflow. Briefly, SH-SY5Y cells were differentiated and then treated with MNP-021 for 48 hours. The MGO stressor was applied for 4 hours. b. Representative images of the toxicity test. Cells were treated with increasing concentrations of MNP-021. Nuclei were labeled with DAPI (blue) and the cytoskeleton using phalloidin (red). Images were acquired with 40x objective. c. Analysis of the ratio between nuclear and cytoplasmic areas to assess MNP-021 toxicity. d. Representative images of SH-SY5Y cells treated with MNP-021 and stressed using MGO. Nuclei were labeled with DAPI (blue) and the cytoskeleton with phalloidin (red). Images were acquired with 40x objective. e. Analysis of the ratio between nuclear and cytoplasmic areas of samples treated with MNP-021, MGO and the combination of the two. Statistical analyses were performed using by One-way ANOVA test. ** ≤0.01. Tukey-Kramer test for multiple comparisons. *: p ≤0.05, **: p ≤0.01, ***: p ≤0.001, ****: p ≤0.0001.

### 3.2 Transcriptomic analysis revealed a transcriptional regulation after MGO and MNP-021 treatments

To define mechanisms underlying MNP-021–mediated protection from MGO-induced damage, we performed RNA-seq to compare gene expression across conditions (Figure 2a) and used pathway enrichment analysis to identify regulated biological programs. After filtering low-count genes, among 24.618 expressed genes, MGO-stressed cells exhibited 754 differentially expressed genes relative to vehicle controls (448 downregulated, 306 upregulated) (Figure 2b). Pathway enrichment analysis indicated that MGO stress upregulated genes primarily involved in apoptosis (Supplementary Figure 1a; Supplementary file 1) and downregulated genes associated with nervous system differentiation and function (Supplementary Figure 1b; Supplementary file 1). Among differentially expressed genes, several were notable for links to glucose handling and the integrated stress response (ISR), including glucokinase (GCK), which promotes glucose utilization/retention^18^, sestrin 2 (SESN2)^19^, phosphoglycerate dehydrogenase (PHGDH), serine biosynthesis/ISR-related (PSAT1)^20^, and mitochondrial anaplerosis/PEP cycling (PCK2)^21^, as shown in Figure 2c. Given the role of SESN2 in mitochondrial integrity and stress responses^19^, this pattern suggests that MGO induces a coordinated stress program with potential perturbation of mitochondrial activity and glucose metabolism. Cells treated with MNP-021 alone showed n= 76 differentially expressed genes versus controls (n = 12 downregulated, n = 64 upregulated) (Figure 2d). Notably, several upregulated genes map to pro-survival mechanisms, including KIT ligand (KITLG), a regulator of proliferation/survival^22^, and AXL receptor tyrosine kinase, a signaling node that activates PI3K/AKT and MAPK pathways and can reduce apoptosis under stress/inflammatory conditions^23,24^ (Figure 2e). Importantly, MNP-021 rescued the expression of MGO-induced GCK, SESN2, PHGDH, PSAT1, and PCK2 toward controls, consistent with normalization of stress-driven metabolic/transcriptional remodeling. Enrichment analysis did not identify statistically significant pathway changes (Supplementary file 1), including apoptosis-related pathways, further supporting a protective profile. MNP-021 also restored physiological expression levels of Patched Domain Containing 4 (PTCHD4) (Figure 2f–g) and Insulin-like Growth Factor Binding Protein 5 (IGFBP5) (Figure 2f-g) in the presence of MGO. PTCHD4 is linked to inhibition of the Sonic Hedgehog (Shh) pathway^25,26^, which regulates neurogenesis, proliferation, and neuronal differentiation and has been linked to protection against oxidative/glucose stress^27,28^. IGFBP5 modulates IGF-1/IGF1R signaling and can influence neuronal survival, proliferation, differentiation, neuroplasticity, and remodeling^29^.

**Figure 2.**
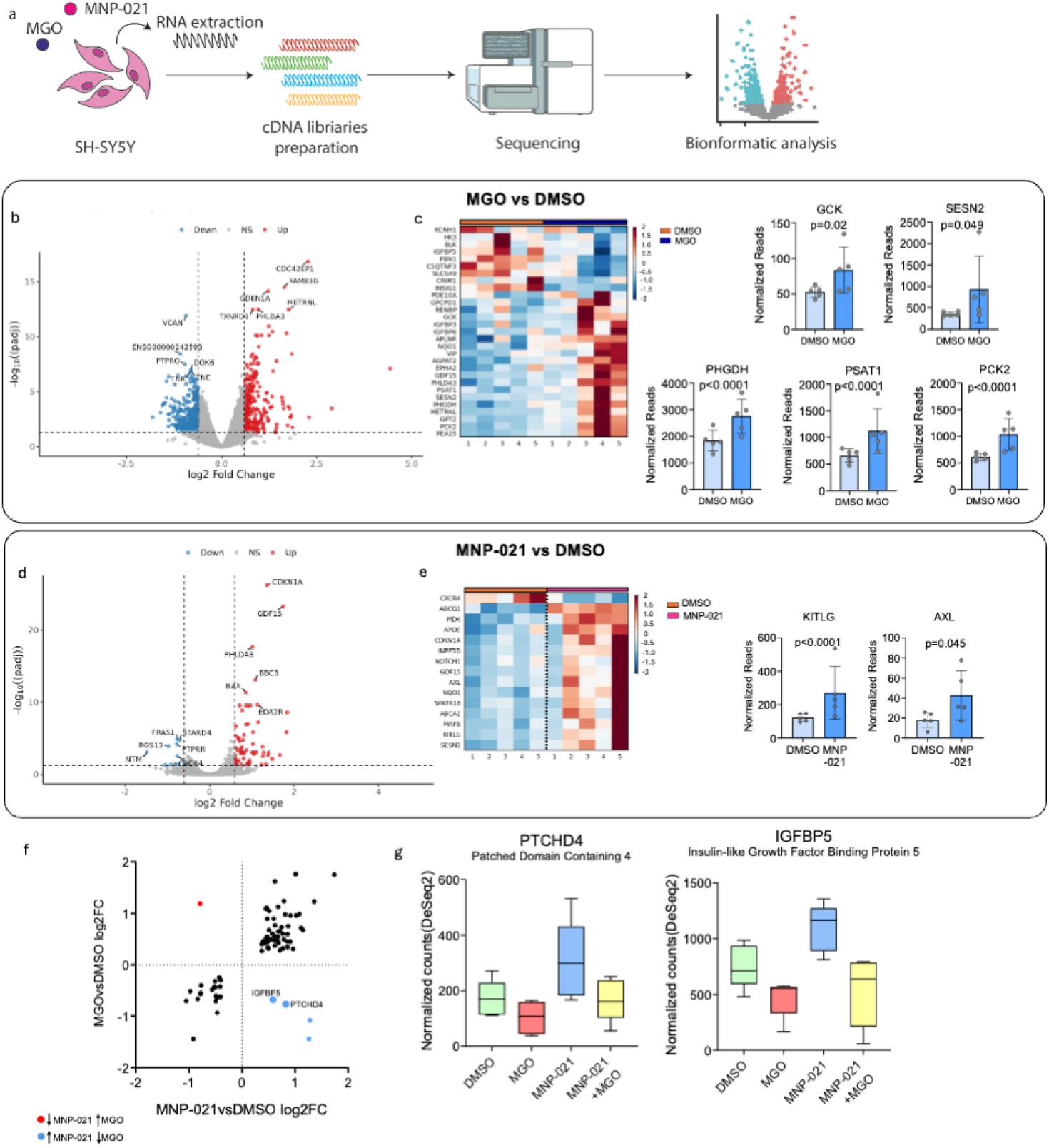
Transcriptomic analysis of SH-SY5Y. a. Experimental Workflow of Transcriptomics analysis performed on SH-SY5Y cell line. b. Volcano plot of n=754 genes differentially expressed in MGO vs CTR. c. Heatmap of n=29 genes differentially expressed in MGO vs CTR and involved in glucose uptake, among which GCK, SESN2, PHGDH, PSAT1, and PCK2 (the most important) are also shown as box plots. d. Volcano plot of n=76 genes differentially expressed in MNP-021 vs CTR. e. Heatmap of n=16 genes differentially expressed in MGO vs CTR and involved in glucose uptake, among which KITL and AXL (the most important) are also shown as box plots. f. Analysis of restored genes showing distribution of n=78 genes differentially expressed in both MGO vs CTR and MNP-021 vs CTR (black, blue and red), as well as the n=5 genes with opposite log_2_FC in MGO vs CTR with respect to MNP-021 vs CTR (blue and red), and n=2 genes not significantly differentially expressed in MGO+MNP-021 vs CTR (blue and bigger). These n=2 genes are also shown in g boxplots for all the experimental conditions.

### 3.3 MNP-021 restores glucose uptake via GLUT transporters

Given the transcriptomic modulation of GCK/SESN2/PHGDH/PSAT1/PCK2, we quantified live-cell glucose uptake by high-content microscopy using 2-NBDG and assessed GLUT1 and GLUT4 localization in differentiated SH-SY5Y cells treated with MNP-021 and/or MGO as above. Following MGO exposure, cells were incubated with 2-NBDG; internalized fluorescence intensity was quantified, with representative images shown in Figure 3a. MGO increased 2-NBDG uptake by ∼25%, whereas MNP-021 counteracts MGO-driven increase of glucose uptake (Figure 3b). To link uptake changes to transporter trafficking, we examined GLUT1/GLUT4 distribution by immunofluorescence. High-content confocal microscopy quantified: (i) nucleus-to-cytoplasm fluorescence ratios; and (ii) membrane vs cytosol compartment signals normalized to total fluorescence, enabling a membrane-to-cytosol redistribution index. GLUT1 staining showed that MGO induced transporter redistribution characterized by membrane accumulation with a reduction in the cytosolic fraction, increasing the membrane-to-cytosol GLUT1 ratio (Figure 3c). MNP-021 counteracted this membrane enrichment and re-equilibrated GLUT1 distribution toward vehicle (Figure 3c). A similar pattern was observed for GLUT4 (Figure 3d): MGO drove marked GLUT4 membrane accumulation at the expense of cytosolic distribution, increasing the membrane-to-cytosol ratio. Quantitative metrics—membrane GLUT4/total GLUT4 (%), cytosolic GLUT4/total GLUT4 (%), and the derived membrane-to-cytosol ratio—confirmed this trafficking shift, which MNP-021 pretreatment fully reversed (Figure 3d).

**Figure 3.**
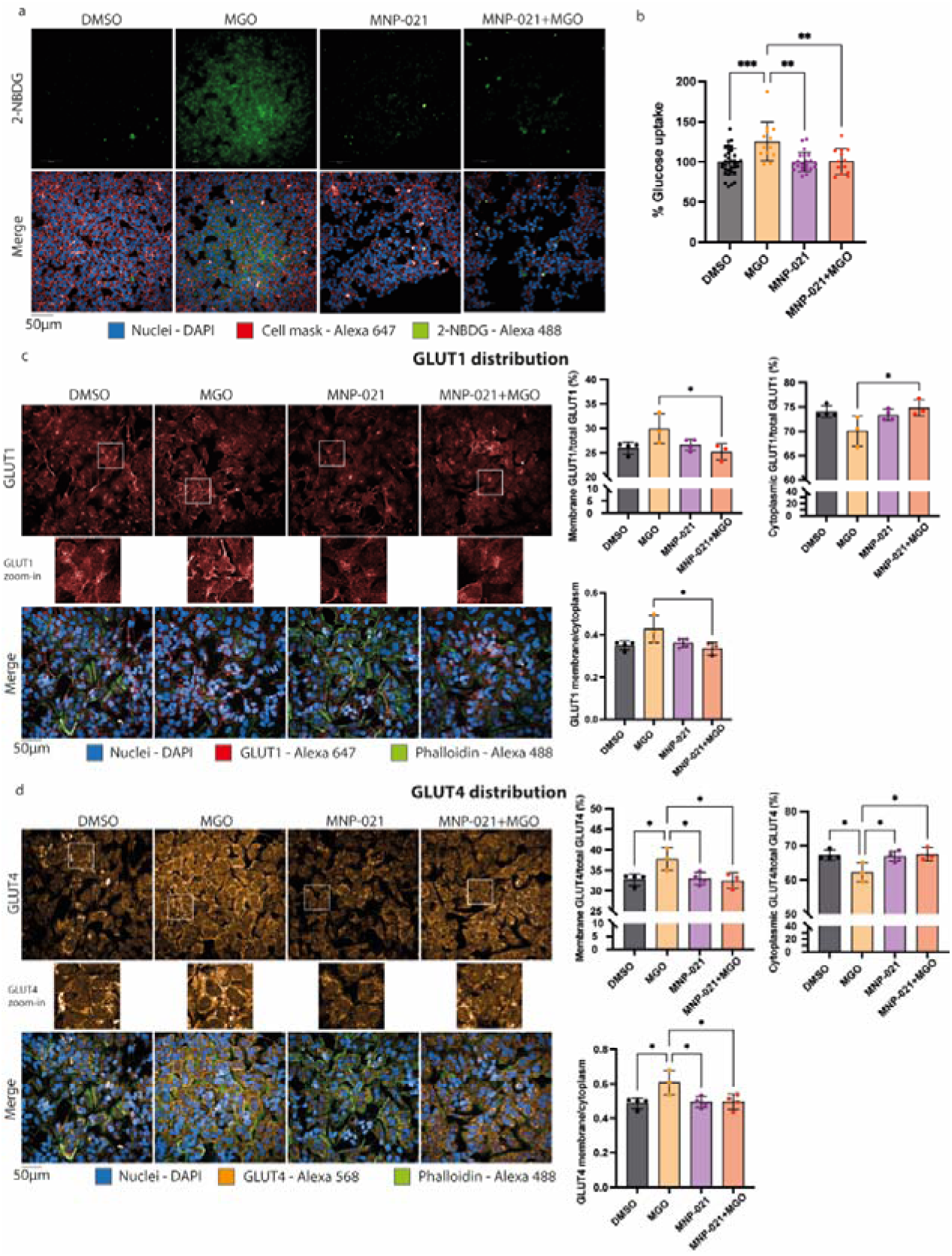
Glucose uptake increased with MGO stressor, but it is restored by MNP-021. a. On the left, there are representative images of glucose uptake. Nuclei were labeled with DAPI (blue), membrane with Cell Mask DeepRed (red), and glucose using 2-NBDG (green). Images were acquired with 40x objective. On the left, statistical analysis of glucose uptake (%) in cells treated with MNP-021, MGO, or a combination of them. Statistical analyses were performed using by One-way ANOVA test. ** ≤0.01. Tukey-Kramer test for multiple comparisons. *: p ≤0.05, **: p ≤0.01, ***: p ≤0.001, ****: p ≤0.0001. b. On the left, there are representative images of GLUT1 distribution in the cells. Nuclei were labeled with DAPI (blue), cytoskeleton with phalloidin (green), and GLUT1 with Alexa647 antibody (red). Images were acquired with 40x objective. On the right, there are statistical analyses of GLUT1 distribution in cells treated with MNP-021, MGO or a combination of them. Statistical analyses were performed by One-way ANOVA test. * ≤0.05. Tukey-Kramer test for multiple comparisons. *: p ≤0.05, **: p ≤0.01, ***: p ≤0.001, ****: p ≤0.0001. c. On the left, there are representative images of GLUT4 distribution in the cells. Nuclei were labeled with DAPI (blue), cytoskeleton with phalloidin (green), and GLUT4 with Alexa568 antibody (orange). Images were acquired with a 40x objective. On the right, there are statistical analyses of GLUT4 distribution in cells treated with MNP-021, MGO, or a combination of them. Statistical analyses were performed using by One-way ANOVA test. * ≤0.05. Tukey-Kramer test for multiple comparisons. *: p ≤0.05, **: p ≤0.01, ***: p ≤0.001, ****: p ≤*0.0001*.

### 3.4 MNP-021 reshapes mitochondrial redox status and ultrastructural features in SH-SY5Y-derived neurons

Prompted by the RNA-seq evidence of stress-responsive metabolic/mitochondrial programs after MGO (including SESN2) and normalization with MNP-021^19^, we assessed mitochondrial structure and bioenergetic function. Differentiated SH-SY5Y cells treated with MNP-021 and exposed to MGO were analyzed for mitochondrial morphology and oxidative readouts using MitoTracker and MitoSOX (Figure 4a–b), followed by Seahorse measuring mitochondrial respiration and ECAR-based glycolysis. High-content imaging revealed a robust redox/morphology signature with MNP-021. Mitochondria in MNP-021–treated cells, in the presence or not of MGO, showed an approximately doubled increase in MitoSOX fluorescence intensity (Figure 4c), consistent with increased mitochondrial ROS-related readouts/oxidative activity. In parallel, MNP-021 increased mitochondrial content: the average number of mitochondria per cell was higher in the MNP-021 and MNP-021 + MGO conditions versus controls (Figure 4c). Morphometric profiling indicated coordinated remodeling, with increased mitochondrial area, length, and roundness under MNP-021 exposure (Figure 4c). To capture an organization beyond size/shape, we quantified mitochondrial texture via SER (structural element response) analysis on MitoTracker signals. Across multiple SER features, MNP-021 (± MGO) significantly reduced index values (Figure 4d), indicating a shift toward a smoother/less structured texture profile compared with DMSO. We then tested whether these imaging phenotypes corresponded to measurable changes in respiratory flux. Seahorse analysis showed that MNP-021 did not significantly modify basal oxygen consumption rate (OCR), ATP-linked respiration, proton leak, FCCP-stimulated maximal OCR, or spare respiratory capacity (Figure 4e). ECAR metrics were likewise not significantly shifted (Figure 4f). Overall, MNP-021 primarily altered mitochondrial abundance/morphology and oxidative readouts without a major detectable reprogramming of bulk respiratory flux. Notably, MGO exposure did not produce the expected impairment of mitochondrial respiration/ETC capacity (Figure 4e–f), likely reflecting the short duration and moderate intensity of the insult compared with paradigms using higher MGO and/or longer exposures (e.g., ∼24 h)^30^.

**Fig 4.**
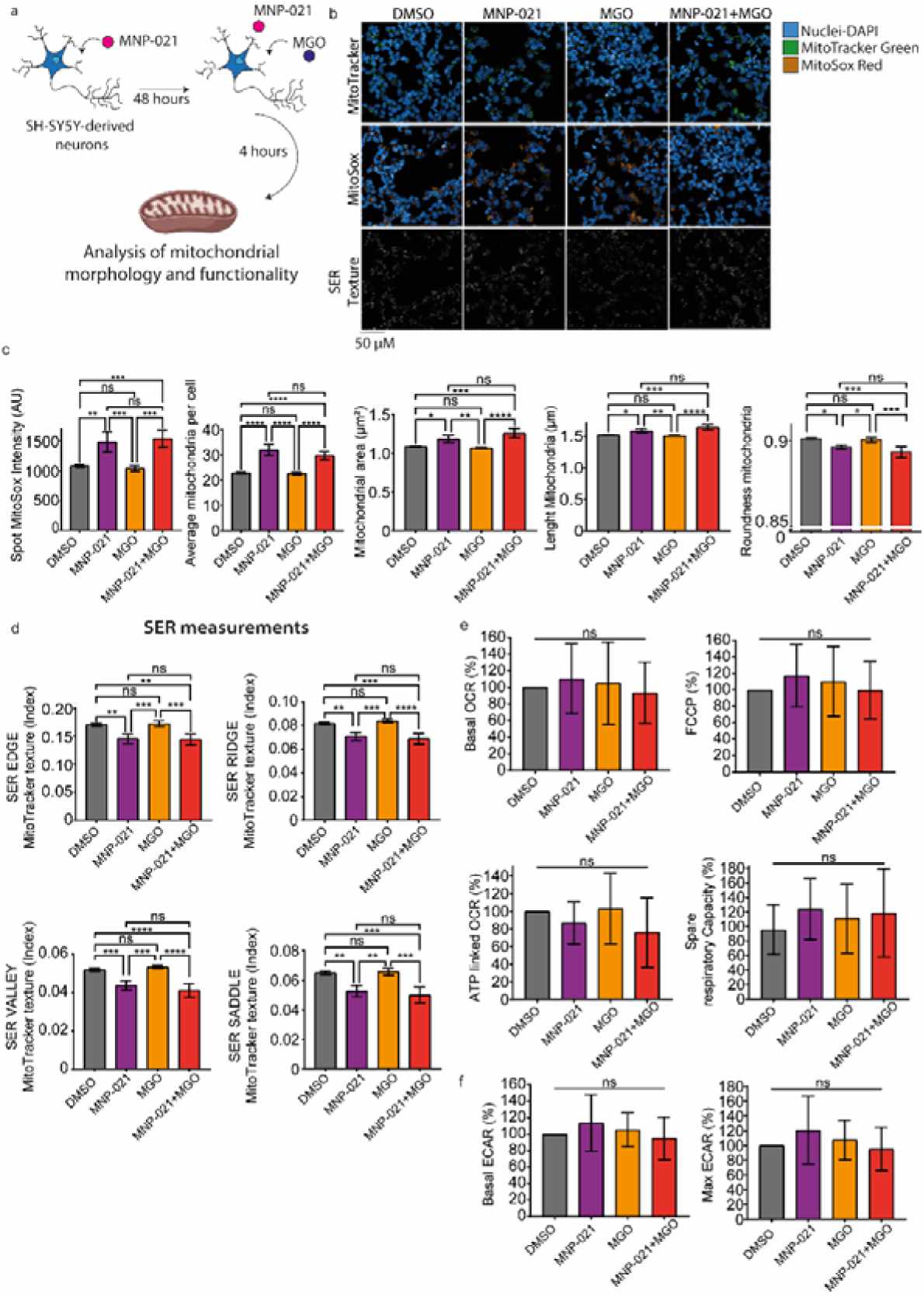
MNP-021 influence on mitochondrial activity and morphology. a. Schematic representation of the experimental workflow. Briefly, SH-SY5Y cells were differentiated and then treated with MNP-021 for 48 hours. The MGO stressor was applied for 4 hours. b. representative images of Mitotracker staining, MitoSox staining, and SER texture. Nuclei were labeled with DAPI (blue), mitochondria and SOX with MitoTracker (green) and MitoSox (red). SER texture was obtained by a specific algorithm in Harmony 5.1 software based on MitoTracker intensity. Images were acquired with a 63x objective. c. Statistical analysis of MitoSox intensity, number of mitochondria, and morphology of mitochondria in cells treated with MNP-021, MGO, or a combination of them. Statistical analyses were performed using by One-way ANOVA test. Tukey-Kramer test for multiple comparisons. *: p ≤0.05, **: p ≤0.01, ***: p ≤0.001, ****: p ≤0.0001. N= 3. d. Statistical analyses of mitochondria SER texture in cells treated with MNP-021, MGO, or a combination of them. Statistical analyses were performed using by One-way ANOVA test. Tukey-Kramer test for multiple comparisons. *: p ≤0.05, **: p ≤0.01, ***: p ≤0.001, ****: p ≤0.0001. N=3. e-f. Statistical analysis of mitochondrial metabolism obtained by SeaHorse assay. Statistical analyses were performed using by One-way ANOVA test. p 0.05, Not-significant. Tukey-Kramer test for multiple comparisons. *: p ≤0.05, **: p ≤0.01, ***: p ≤0.001, ****: p ≤0.0001. N=8.

### 3.5 MNP-021 attenuates MGO-driven glycolytic stress responses

To resolve treatment-related changes in acidification dynamics not captured by steady-state ECAR, we performed a glycolytic stress assay (Figure 5). ECAR was quantified across sequential phases (FRB/basal glycolysis; glucose + oligomycin to elicit compensatory glycolysis; and 2-DG + glucose to estimate non-glycolytic acidification), allowing decomposition into basal, compensatory, and non-glycolytic components (Figure 5a–b). MGO significantly increased acidification during the oligomycin phase, consistent with enhanced glycolytic compensation under stress (Figure 5b–c). MNP-021 alone maintained ECAR/acidification profiles comparable to DMSO controls (Figure 5b), indicating no intrinsic pro-glycolytic perturbation. Importantly, MNP-021 + MGO co-treatment blunted the MGO-induced acidification rise (Figure 5b–c), supporting protection against stress-evoked glycolytic overactivation. As expected, GOX produced the highest acidification responses, confirming assay sensitivity (Supplementary Figure 2; Supplementary Table 4). Untreated cells showed low acidification across phases, consistent with minimal basal metabolic activation (Figure 5c–d).

**Figure 5:**
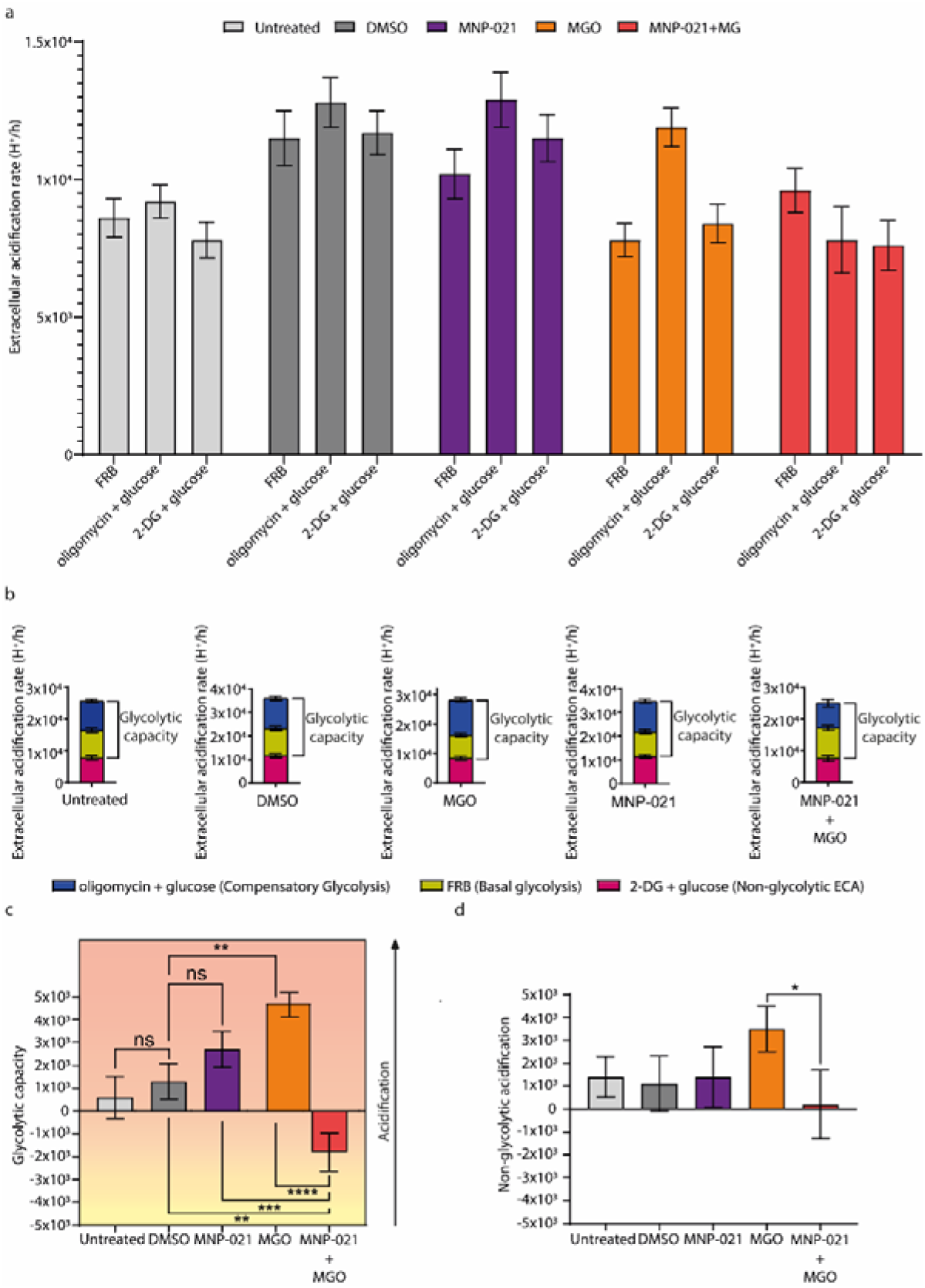
MNP-021 + MGO co-treatment significantly attenuated MGO-induced acidification. a. Graphical representation of the Extracellular acidification rate in RFB, oligomycin + glucose, and 2-DG + glucose phases assay for each treatment. b. Individual representation of RFB (basal glycolysis), oligomycin + glucose (Compensatory glycolysis), and 2-DG + glucose (Non-glycolytic ECA) per treatment. c. Glycolytic capacity representation, indicating the acidification rate in the sample in comparison to the baseline. Statistical analyses were performed using by One-way ANOVA test. Tukey-Kramer test for multiple comparisons. *: p ≤0.05, **: p ≤0.01, ***: p ≤0.001, ****: p ≤0.0001. N= 3. d. Non-glycolytic acidification representation. Statistical analyses were performed using by One-way ANOVA test. Tukey-Kramer test for multiple comparisons. *: p ≤0.05, **: p ≤0.01, ***: p ≤0.001, ****: p ≤0.0001. N=3.

### 3.6 MNP-021 restores pAKT/pERK/CREB signaling

Based on transcriptomic evidence consistent with modulation of programs linked to neurogenesis, proliferation, and differentiation, we tested whether MNP-021 engages canonical pro-survival/pro-plasticity signaling cascades in neurons. We focused on AKT, ERK1/2, and CREB, given their roles in neuronal survival, differentiation, and plasticity^31–34^. Differentiated SH-SY5Y cells were analyzed by western blot to quantify phosphorylated (active) proteins relative to total levels. AKT activation (pAKT/AKT) showed that MGO caused a ∼20% decrease in basal pAKT (non-significant), whereas MNP-021 significantly increased pAKT in the presence or not of MGO (Figure 6a), supporting robust engagement of AKT signaling. The administration of insulin increased pAKT by ∼9-times, while MGO reduced insulin-driven AKT activation by ∼2-fold. Notably, MNP-021 restored and further enhanced AKT phosphorylation in the presence of MGO during insulin stimulation (Figure 6b). ERK1/2 displayed a concordant pattern: in basal conditions, MNP-021 increased pERK1/2 to ∼3-times above controls ± MGO (Figure 6c). Under insulin stimulation, ERK1/2 activation was strongly induced, MGO reduced pERK1/2 by ∼2-fold versus insulin controls, and MNP-021 restored and further increased ERK1/2 activation (Figure 6d). Time-course analysis (30–180 min; Figure 6e) showed sustained ERK1/2 suppression with MGO (∼25% reduction early onward), while MNP-021 induced rapid pERK1/2 elevation (from 15 min) and maintained increased activation even under MGO challenge (Figure 6e). Nuclear fractionation supported functional pathway engagement: MGO did not reduce nuclear pERK1/2 significantly, whereas MNP-021 elevated nuclear pERK1/2 in the presence or not of MGO (Figure 6f). Given CREB as a downstream integrator of AKT and ERK1/2 signaling in neurons^34^, we quantified CREB activation kinetics. MNP-021 induced a rapid and sustained pCREB increase detectable within 15 min (∼16%) and peaking around 60 min (additional ∼35%) relative to untreated cells (Figure 6g). Nuclear extracts showed increased phosphorylated CREB accumulation with MNP-021 (∼30% rise in pCREB signal) (Figure 6h). Collectively, MNP-021 activates AKT and ERK1/2 signaling and promotes downstream CREB phosphorylation, including under MGO stress and during insulin stimulation, supporting preservation of neuronal signaling competence and pro-survival/pro-plasticity programs^31–34^.

**Figure 6.**
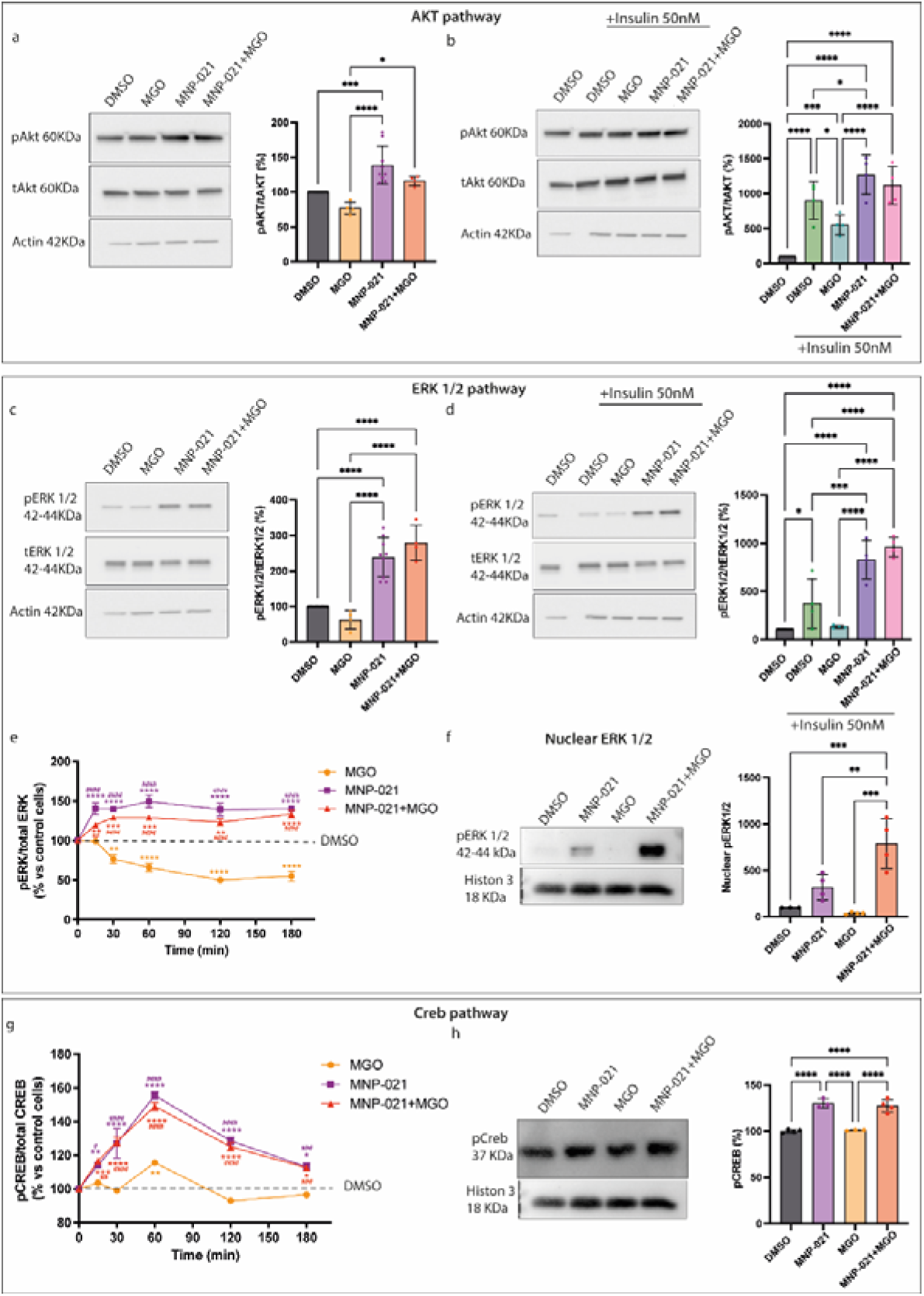
MNP-021 activates AKT, ERK1/2, and CREB pathways. a. Western blot representative images and analysis of the percentage of pAKT on the total AKT (tAKT) content in differentiated SH-SY5Y cells (a) treated or not with MNP-021, MGO, or the combination of the two, (b) and in the presence or not of insulin. Statistical analyses were performed using by One-way ANOVA test. Tukey-Kramer test for multiple comparisons. *: p ≤0.05, **: p ≤0.01, ***: p ≤0.001, ****: p ≤0.0001. c,d. Western blot images and analysis of the percentage of pERK1/2 on total ERK1/2 (tERK1/2) content in SH-SY5Y cells (c) treated with MNP-021, MGO, or the combination of the two, (d) and in the presence or not of insulin. Statistical analyses were performed using by One-way ANOVA test. Tukey-Kramer test for multiple comparisons. *: p ≤0.05, **: p ≤0.01, ***: p ≤0.001, ****: p ≤0.0001. e. Kinetic analysis of ERK1/2 activation measured by immunoenzymatic assay. Differentiated SH-SY5Y were then treated and analyzed at different time points (0,30, 60, 90, 120, 150, 180 minutes). The data was expressed as pERK1/2 on tERK1/2 content. Statistical analyses were performed using by Two-way ANOVA test. Tukey-Kramer test for multiple comparisons. *: p ≤0.05, **: p ≤0.01, ***: p ≤0.001, ****: p ≤0.0001. N= 4. *: indicates the statistical significance among different time points of the same treatment, #: indicates the statistical significance among different treatments at the same time points. f. Western blot showing pERK1/2 content in the nuclear fraction. Protein content was normalized on a nuclear housekeeping protein, Histone-3 protein. Statistical analyses were performed using by One-way ANOVA test. Tukey-Kramer test for multiple comparisons. *: p ≤0.05, **: p ≤0.01, ***: p ≤0.001, ****: p ≤0.0001. g. Kinetic analysis of CREB activation measured by immunoenzymatic assay. Differentiated SH-SY5Y were then treated with MNP-021 and MGO, and analyzed at different time points (0,30, 60, 90, 120, 150, 180 minutes). The data was expressed as pCREB on total CREB content. Statistical analyses were performed using by Two-way ANOVA test. Tukey-Kramer test for multiple comparisons. *: p ≤0.05, **: p ≤0.01, ***: p ≤0.001, ****: p ≤0.0001. N= 4. *: indicates the statistical significance among different time points of the same treatment, #: indicates the statistical significance among different treatments at the same time points. h. Western blot analysis of the nuclear content of pCREB, normalized on the Histon 3 protein content. The data were shown as the mean ± SEM of individual experiments. Statistical analyses were performed using by One-way ANOVA test. Tukey-Kramer test for multiple comparisons. *: p ≤0.05, **: p ≤0.01, ***: p ≤0.001, ****: p ≤0.0001.

### 3.7 MNP-021 prevents microglia inflammatory response

In diabetes, chronic hyperglycemia and metabolic dysregulation contribute to sustained microglia activation and a pro-inflammatory phenotype with elevated cytokines (e.g., TNF-α, IL-6), accelerating diabetic neuropathy and cognitive decline^35^. Before testing anti-inflammatory effects, we assessed MNP-021 cytotoxicity in HMC3 microglia cells: 24h treatment across concentrations did not reduce viability by MTT, indicating the absence of cytotoxicity even at the highest doses (Figure 7a).

**Figure 7.**
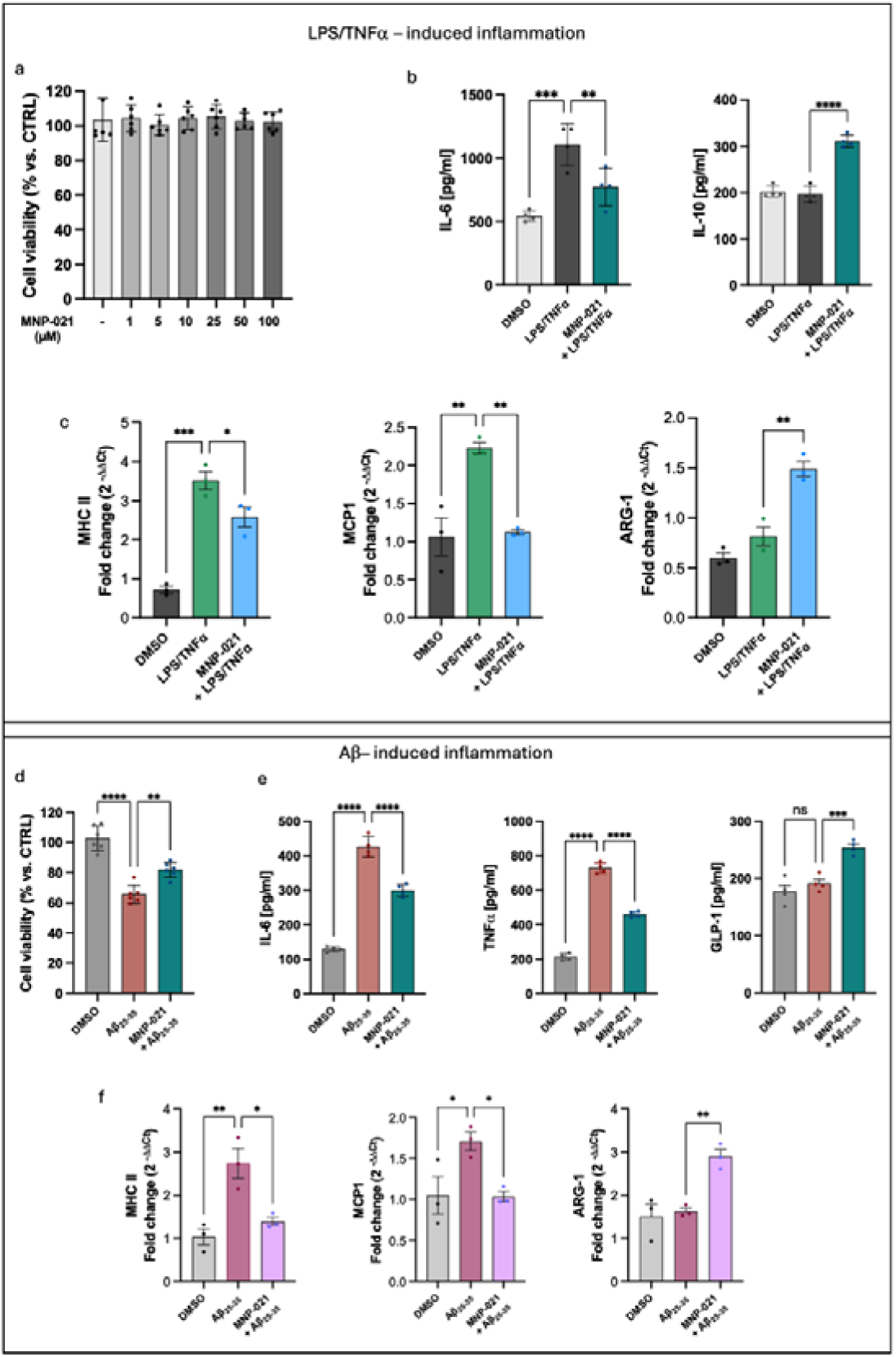
Anti-inflammatory effects of MNP-021 in HMC3 microglial cells. a. Treatment with MNP-021 (1– 100 μM, 24 h) did not affect HMC3 cell viability. b. MNP-021 pre-treatment significantly reduced LPS/TNF-α-induced IL-6 release and increased IL-10 secretion. c. LPS/TNF-α stimulation upregulated the pro-inflammatory markers MHC class II and MCP1, while ARG1 expression was unchanged; MNP-021 prevented the induction of pro-inflammatory markers and increased ARG1 expression. d. MNP-021 partially protected HMC3 cells from Aβ□□□–□□□induced cytotoxicity. e. Aβ□□□–□□□increased IL-6 and TNF-α release without affecting IL-10, while MNP-021 pre-treatment attenuated pro-inflammatory cytokines and enhanced IL-10 secretion. (f) Consistently, MNP-021 prevented Aβ□□□–□□□-induced upregulation of MHC class II and MCP1 and significantly increased ARG1 expression. Statistical analyses were performed by One-way ANOVA test followed by Dunnett’s test for multiple comparisons. *: p ≤0.05, **: p ≤0.01, ***: p

As reported, stimulation of HMC3 cells with LPS + TNF-α for 24 h induced a robust inflammatory response with increased IL-6 release and no significant IL-10 change versus unstimulated controls^15,36^. With 10 µM MNP-021 pretreatment for 1 h before LPS/TNF-α and 24 h incubation, ELISA showed that MNP-021 prevented the LPS/TNF-α–induced IL-6 increase and significantly promoted IL-10 secretion versus LPS/TNF-α alone (Figure 7b). To profile microglial activation state, qPCR assessed MHC II and MCP1 (pro-inflammatory/M1-like markers)^12,37^ and ARG1 (M2-like marker)^13^. LPS/TNF-α increased MHC II and MCP1, while ARG1 remained comparable to controls; MNP-021 prevented MHC II and MCP1 upregulation and induced an ∼1.9-fold increase in ARG1 (Figure 7c), consistent with attenuation of pro-inflammatory activation and a shift toward an anti-inflammatory program. We next used an Aβ-based neuroinflammation model in HMC3 cells, since Aβ_(25–35)_ induces microglial activation and cytotoxicity^38,39^. Aβ reduced viability by ∼35% and increased TNF-α and IL-6 secretion, without changing IL-10 versus control (Figure 7d). MNP-021 pretreatment attenuated the Aβ-induced TNF-α and IL-6 increases though remaining above control and significantly increased IL-10 in Aβ-exposed cells (Figure 7e). qPCR confirmed that Aβ upregulated MHC II and MCP1 with no ARG1 change (Figure 7f), while MNP-021 prevented MHC II/MCP1 upregulation and increased ARG1, reinforcing anti-inflammatory reprogramming (Figure 7f). Finally, we evaluated whether MNP-021 modulates GLP-1 release in HMC3 cells, given neuroprotective/anti-inflammatory roles of GLP-1 signaling^40^. GLP-1 ELISA in media from MNP-021–pretreated HMC3 cells stimulated with LPS/TNF-α showed no significant increase; concentrations remained very low (≤10 pg/mL) (Supplementary figure 3). Although GLP-1 expression has been reported in HMC3 cells^41^, endogenous GLP-1 production is likely not a relevant mechanism here (Supplementary Figure 3), suggesting that MNP-021 protection is mediated by alternative pathways.

## 4. Discussion

In this study, we position the TRPA1-directed small molecule MNP-021 as a multi-compartment modulator of neuroglial damage in experimental contexts that capture key pathobiological nodes of diabetes-associated neurodegeneration, where MGO and related dicarbonyl stressors link hyperglycemia to mitochondrial impairment, insulin resistance–like signaling defects, and maladaptive neuroinflammation^5,9,42^. In differentiated SH-SY5Y neurons subjected to acute MGO challenge, MNP-021 preserved cellular architecture, restrained the amplitude of stress-induced transcriptional remodeling, normalized glucose handling together with GLUT1/GLUT4 distribution, buffered glycolytic stress compensation, restored pro-survival kinase signaling such as AKT, ERK1/2 and CREB.

In addition, we found that in human HMC3 microglia cells, MNP-021 shifted inflammatory activation toward a less injurious, more pro-resolving profile. These evidences together outline a coherent “stress-rheostat” phenotype rather than a single-pathway antioxidant effect. RNA-seq profiling identified 754 MGO-deregulated transcripts, of which 448 were found downregulated while 306 up-regulated. Overall, transcriptomic analyses revealed a broad integrated stress response caused by MGO administration, with coordinated upregulation of glycolytic control and mitochondrial-related stress responses, consistent with omics datasets showing that MGO perturbs neuronal metabolic networks, proteostasis, and stress signaling in SH-SY5Y models^9,43^. Importantly, these differences are selectively blunted by MNP-021 in cells under MGO-induced stress, implying normalization of adaptive-to-injurious reprogramming without basal transcriptome perturbation. A central mechanistic advance of this work is the coupling of live-cell functional glucose uptake (2-NBDG) to compartment-resolved assessment of GLUT1/GLUT4 organization. Of note, MGO increased intracellular 2-NBDG accumulation (∼25%), whereas MNP-021 pre-treatment counteracts MGO-induced increase in glucose uptake. This evidence aligns with the observed preservation of GLUT1/GLUT4 spatial distribution.

While the literature reports variable directions of MGO-induced glucose-uptake changes depending on dose, timing, and differentiation status, successful interventions consistently converge on restoring metabolic competence; sulforaphane, for instance, mitigates MGO toxicity while enhancing glyoxalase/antioxidant defenses^44^. The transporter-centric layer is particularly relevant because neuronal GLUT4 trafficking is dynamically regulated by insulin/leptin via PI3K-dependent pathways, directly linking trophic signaling to substrate availability and neuronal insulin sensitivity^45^. Compared with paradigms that mainly rescue downstream redox/mitochondrial injury (e.g., carnosic acid via PI3K/Akt–Nrf2–GSH; liraglutide via energy-metabolism rebalancing)^46,47^. MNP-021 aligns neuroprotection with transporter-level homeostasis, consistent with a proximal “metabolic stabilizer” effect at the signaling–trafficking interface.

Mitochondrial profiling further supports this interpretation. Across the short exposure window, MGO induced only modest changes in bioenergetic readouts, whereas high-content imaging revealed that MNP-021 increased mitochondrial abundance while preserving a predominantly elongated/filamentous morphology and reshaping mitochondrial redox-associated signals. These effects on mitochondria were not related to major shifts in OCR/ECAR parameters in this exposure window, suggesting stabilization of structure, signaling coupling and stress readiness rather than overt reprogramming of basal respiration. This structural protection occurred alongside increased MitoSOX signal with MNP-021, alone or with MGO, arguing against a simplistic ROS-scavenger mechanism and instead aligning with controlled ROS signaling/mitohormesis concepts in which modest mitochondrial superoxide can reinforce resilience programs^46^. This is compatible with the broader MGO literature, where neuronal injury often features mitochondrial depolarization, in which ATP loss and ROS rise are commonly described. In this context, protection can be achieved via AGEs/RAGE/NF-κB modulation, Nrf2/GSH engagement, and related mitochondrial-defense axes (e.g., myricitrin; naringenin)^48,49^. Mechanistically, stress-inducible guardians such as SESN2 can preserve mitochondrial integrity and mitophagy competence under diverse stressors, motivating direct interrogation of SESN2/mitophagy nodes downstream of MNP-021^50^. Consistent with a coordinated repair program, immunoblotting showed that MNP-021 increased pAKT(Ser473) and pERK1/2(Thr202/Tyr204) and enhanced nuclear pCREB(Ser133), counteracting MGO-induced signaling fragility and fitting with the concept of “dicarbonyl-driven insulin resistance” at the IRS–PI3K interface described in neuronal/metabolic models, including ROS-independent impairment of IRS-1–proximal signaling^51^. Given that nuclear AKT integrates neuronal survival and transcriptional programs, and that ERK1/2–CREB coupling supports plasticity and protection-associated gene expression^32,34^, simultaneous restoration of AKT/ERK/CREB together with glucose-transport and mitochondrial-architecture phenotypes suggests that MNP-021 helps re-establish signaling-to-metabolism homeostasis during dicarbonyl stressors. Notably, several neuroprotection paradigms show causal dependence on PI3K/AKT/CREB (e.g., rifampicin protection against rotenone is PI3K-sensitive; ginsenoside Rb1 and carnosic acid converge on PI3K/AKT-linked defenses), reinforcing pathway plausibility^52,53^. Extending beyond neurons, MNP-021 also modulated microglial inflammatory tone. Microglia can amplify neurodegeneration or support repair depending on activation state^54^. Although HMC3 identity has been debated, this cell line remains widely used for human microglia-like screening and inflammatory phenotyping^55,56^. In our setting, LPS/β-amyloid drove a canonical pro-inflammatory program (increased IL-6, MCP-1, MHC-II; reduced IL-10; altered phagocytosis) consistent with prior work linking IL-10 loss to an M1-prone phenotype^57^. MNP-021 partially reversed this state by reducing IL-6/MCP-1 and in parallel increasing IL-10 and Arg1. Moreover, the drug preserved viability and phagocytic competence, supporting qualitative reprogramming rather than global immunosuppression. IL-10/Arg1-enriched microglial/macrophage states have been linked to improved outcomes in experimental injury contexts^58^. A key conceptual tension is that MGO can activate TRPA1, including from the intracellular side, contributing to nociceptor sensitization and mechanisms relevant to metabolic neuropathies^7,59^. Therefore, TRPA1-directed modulation might appear paradoxical. Our data support a resolution in which mode and context of TRPA1 engagement are decisive: non-electrophilic modulation may tune channel microdomain signaling and downstream adaptation rather than promoting sustained Ca²□overload. Indeed, TRPA1 outputs can differ across cellular contexts according to Ca²□buffering, redox tone, and effector wiring^60^. In this framework, MNP-021 acts less as a blunt “TRPA1 activator” and more as a calibrated stress rheostat that decouples MGO exposure from deleterious trajectories while preserving or amplifying protective programs. This hypothesis now requires definitive target-validation by creating TRPA1 loss-of-function/CRISPR or pharmacologic epistasis in both neurons and microglia.

Translationally, MNP-021 is attractive because it targets convergent mechanisms linking dysmetabolism to neurodegeneration rather than a single downstream effector, potentially complementing glucose-lowering therapies that incompletely address cognitive decline risk^5,42^.

The present work is limited by using immortalized cell lines in which an acute stress paradigm may not capture chronic adaptation, by the absence of genetic on-target validation and the lack of *in vivo* confirmation of brain exposure, with region-specific efficacy and long-term safety.

In conclusion, MNP-021 displays a distinctive, multi-layer neuroprotective signature that integrates transcriptomic restraint, transporter-level glucose handling, mitochondrial architectural resilience, reinstatement of AKT/ERK/CREB signaling, and microglial reprogramming. If these coordinated effects translate beyond *in vitro* models, TRPA1-tuned small-molecule scaffolds such as MNP-021 could represent a new strategy to intercept the MGO–TRPA1– mitochondria–microglia axis early, with potential relevance to diabetic neuropathy, cognitive impairment, and broader neurodegenerative trajectories in metabolic disease.

## Declaration of competing interest

Giuseppe Daniele and Mauro Pineschi are inventors on patent applications covering MNP-021 and related compounds (PCT filed August 2023; published as WO2025/032536; with related patent family WO2018/220542; national/regional phases in IT/EP/US). The other authors declare no competing interests.

## Authorship contribution statement

Alessandro De Carli, Carolina Filipponi, Michele Lai: Data curation, Formal analysis, Investigation, Visualization, Writing original draft. Beatrice Polini, Emy Basso, Giuseppina Emanuela Grieco, Guido Sebastiani, Maria Grazia Chiellini, Simona Daniele, Martina Contestabile: Investigation, Methodology, Writing, review and editing. Francesco Dotta, Angela Dardano, Francesca Sardelli: review. Veronica Sancho-Bornez, Fabio Filippini: Investigation, Data curation. Giuseppe Daniele: Conceptualization, Methodology, Project administration, Supervision, Funding acquisition, Writing original draft, review and editing. Mauro Pineschi: Writing, review and editing.

## Acknowledgments

The authors thank Giusy Zappulla, Giorgia Bray, and the technical staff of the participating laboratories for their valuable support with experimental procedures, sample handling, and day-to-day laboratory operations. This work was supported by institutional research funds from the University of Pisa and by the University of Pisa-Bando Dimostratori Tecnologici 2022 (Decreto Rettorale D.R. 1286/2022, 22 July 2022; award list published by D.R. 53/2023, 12 January 2023)

## Data availability

The data sets generated during this study are available from the corresponding author upon reasonable request.

## Highlights

- A non-electrophilic TRPA1 modulator MNP-021 protects neurons from methylglyoxal stress
- MNP-021 normalizes glucose uptake and GLUT1/GLUT4 trafficking under stress
- It preserves mitochondrial architecture and blunts stress-evoked glycolysis
- It restores AKT/ERK/CREB signaling and insulin responsiveness during glucotoxicity
- It reprograms microglia toward pro-resolving IL-10/ARG1 responses

