## Supplementary material for "Unconventional Small Molecule MNP-021 Protects Neuronal and Glial Function from Diabetes-Associated Glucotoxicity and Neuroinflammation"

**Supplementary material and methods**

***Analyses of High-content confocal images***

Images were processed using Harmony 5.1 Software (Revvity, Waltham, MA, USA) following these building blocks:

*For the morphological analysis:* Find Nuclei > Find Cytoplasm > Calculate morphology properties > Calculate ratio Area Nuclei/Area Cytoplasm.

*For the analysis of GLUT localization:* Find Nuclei > Find Cytoplasm > Find Cell Region (Cell membrane) > Find Spots in the cytoplasm > Calculate Spot intensity properties > Select Spot population (based on fluorescence intensity) > Find Spots on the membrane > Calculate Spot intensity properties > Select Spot population (based on fluorescence intensity).

*For the glucose uptake:* Find Nuclei > Find Cytoplasm > Calculate intensity properties in the cytoplasm.

*For MitoSOX fluorescence:* Find Nuclei > Find Cytoplasm > Find Spots in the cytoplasm > Calculate Spots intensity properties.

*For MitoTracker*: Find Nuclei > Find Cytoplasm > Find Spots in the cytoplasm > Find Spot Morphology > Find Spot Texture (SER measurements).

***Separation of the cell components: cytosol, membrane, and nucleus***

To separate cytosolic, membrane, and nuclear fractions, we employed a differential centrifugation protocol. Cells were homogenized on ice using a Potter-Elvehjem homogenizer with ~30 strokes to ensure mechanical disruption, in lysis buffer (10 mM TRIS, pH 7.4, 5 mM EDTA, 5 µg/ml benzamidine, and protease inhibitors cocktail), followed by centrifugation at 10,000 × g for 10 min at 4ºC. The pellet, containing the nuclear fraction, was separated from the supernatant, containing cytosolic and membrane fractions. The cytosolic and membrane fraction supernatant was centrifuged at 40,000 × g for 20 minutes at 4°C to split the cytosolic fraction (supernatant) from the membrane fraction (pellet). Cytosolic and Nuclear fractions were then processed for Western blot analysis following standard lysate preparation protocols.

***Lysate samples preparation and Western blot***

Cell pellets were lysed in RIPA buffer (9.1 mM NaH2PO4, 1.7 mM Na2HPO4, 150 mM NaCl, pH 7.4, 0.5% sodium deoxycholate, 1% Nonidet P-40, 0.1% SDS, and a protease-inhibitor cocktail) by sonication (amplitude 35%, for 3 times for 30 seconds each). Then the samples were lysed for 2h at 4ºC.

Following protein quantification, equal amounts of protein (~40 µg) were loaded onto 7.5% precast SDS-PAGE gels. After electrophoresis, proteins were transferred to PVDF membranes (Bio-Rad, Milan, Italy). Subsequently, membranes were incubated overnight at 4°C with primary antibodies diluted in 5% milk/TBS-T solution (Supplementary Table 1 for antibody details). The next day, membranes were incubated with the appropriate secondary antibodies for 2 hours at RT with gentle shaking. Densitometric analysis of immunoreactive bands was performed using Image Lab Software (Bio-Rad, Milan, Italy). Results were expressed as a percentage relative to the control group. Western blot analyses were carried out using the ChemiDoc XRS+ Gel Imaging System (Bio-Rad, Milan, Italy), with stain-free total protein normalization. This approach considers the total lane protein signal. Consequently, the use of housekeeping proteins was not required. However, to ensure proper fractionation and validation of component separation, GAPDH was used as a control for the cytosolic and membrane fractions, while histone H3 was used as a control for the nuclear fraction.

**Supplementary Table 1.** Primary antibodies used in the Western Blot analysis.

| Protein | Catalog number, supplier | Dilution |
|---|---|---|
| **GLUT1** | #12939, Cell Signaling, USA | 1:500 |
| **GLUT4** | #2213, Cell Signaling, USA | 1:500 |
| **p-CREB** | #9197, Cell Signaling, USA | 1:500 |
| **p-ERK** | sc-7983, Santa Cruz, USA | 1:200 |
| **GAPDH** | G9545, Sigma-Aldrich, USA | 1:1000 |
| **Histone $H_3$** | sc-517576, Santa Cruz, Milan, Italy | 1:200 |

##### *Kinetic analysis by immunoenzymatic assay*

**Supplementary Table 2.** Primary antibodies used in the immunoenzymatic assay analysis.

| Protein | Catalog number, supplier | Dilution |
|---|---|---|
| **p-AKT** | sc-7985-R, Santa Cruz, USA | 1:300 |
| **p-CREB** | #9197, Cell Signaling, USA | 1:300 |
| **p-ERK** | sc-7983, Santa Cruz, USA | 1:300 |
| **Total AKT** | sc-81434, Santa Cruz, USA | 1:300 |
| **Total ERK** | sc-514302, Santa Cruz, USA | 1:300 |

##### *Cell density normalization*

To normalize cell density and confirm equal seeding across wells, after the end of the experiment, wells were stained with crystal violet. After completion of the TMB assay and absorbance measurement, cells in the reference plate were incubated with 4% formaldehyde at RT for 20 minutes and washed twice with PBS 1×. After, cells were stained with 0.5% crystal violet solution for 20 minutes. The bound dye was then solubilized with 1% SDS solution diluted in PBS and incubated for 1 hour on a shaker plate. Then, the violet solution was transferred to a new 96-well ELISA plate, and the absorbance was measured at 590 nm using a microplate reader.

At least two independent experiments were conducted, each with triplicate samples, and the median values were used for analysis. Results were expressed as a percentage relative to the control group.

##### *Cell viability assay (MTT) on HMC3 cells*

Cell viability was assessed using the 3-(4,5-dimethylthiazol-2-yl)-2,5-diphenyltetrazolium bromide (MTT) assay. Briefly, HMC3 cells were seeded in 96-well plates and treated with increasing concentrations of MNP-021 (1–100 μM) for 24 h at 37 °C. At the end of the treatment, MTT solution (5 mg/mL) was added to each well and incubated for 2 h at 37 °C. The culture medium was then removed, and the resulting formazan crystals were solubilized by adding 50 μL of DMSO per well. After a 10-min incubation at 37 °C, absorbance was measured at 540 nm using an automated microplate reader (BIO-TEK, Winooski, VT, USA). Cell viability was expressed as a percentage relative to vehicle-treated control cells.

The same experimental procedure was applied to evaluate the cytotoxic effects induced by $A\beta_{25–35}$ (10 μM, 24 h) in the absence or presence of MNP-021.

##### *RNA extraction and RT-qPCR analysis on HMC3 cells*

**Supplementary Table 3.** Primer sequences used in qRT-PCR on HMC3 cells.

| Gene | Forward primer | Reverse primer |
|---|---|---|
| **MHC II** | AGCTGTGGACAAAGCCAACCTG | CTCTCAGTTCCACAGGGCTGTT |
| **MCP1** | AGAATCACCAGCAGCAAGTGTCC | TCCTGAACCCACTTCTGCTTGG |
| **ARG-1** | TCATCTGGGTGGATGCTCACAC | GAGAATCCTGGCACATCGGGAA |
| **GAPDH** | GTCTCCTCTGACTTCAACAGCG | ACCACCCTGTTGCTGTAGCCAA |

##### *GLP-1 quantification by ELISA*

Glucagon-like peptide-1 (GLP-1) levels were quantified using a commercial human GLP-1 ELISA kit (Cat. #18381624, Merk Millipore), following the manufacturer's instructions. Absorbance was measured at the recommended wavelength using a microplate reader, and GLP-1 concentrations were calculated based on the standard curve. The sensitivity of the assay allowed detection of GLP-1 levels in the low pg/mL range.

### Supplementary results

#### *Transcriptomic analysis revealed a transcriptional regulation after MGO and MNP-021 treatments*

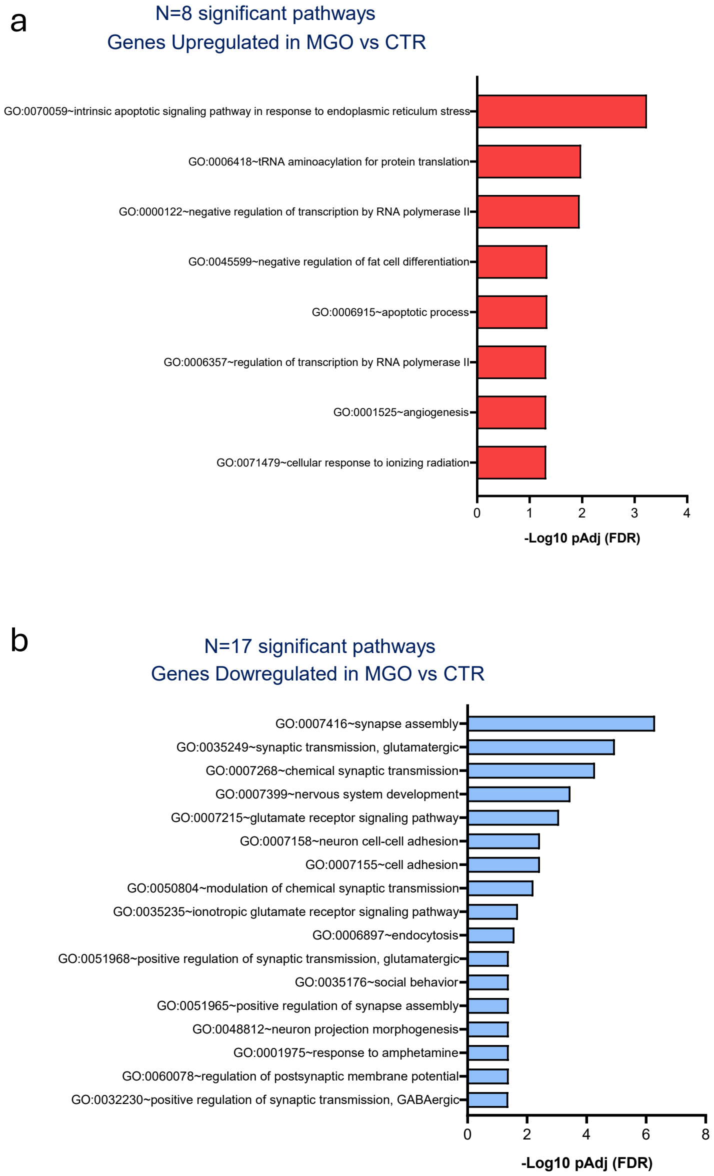


**Supplementary Figure 1**. Pathways enrichment analysis of (a) genes upregulated in MGO vs CTR and of (b) genes downregulated in MGO vs CTR.

#### *MNP-021 attenuates MGO-driven glycolytic stress responses*

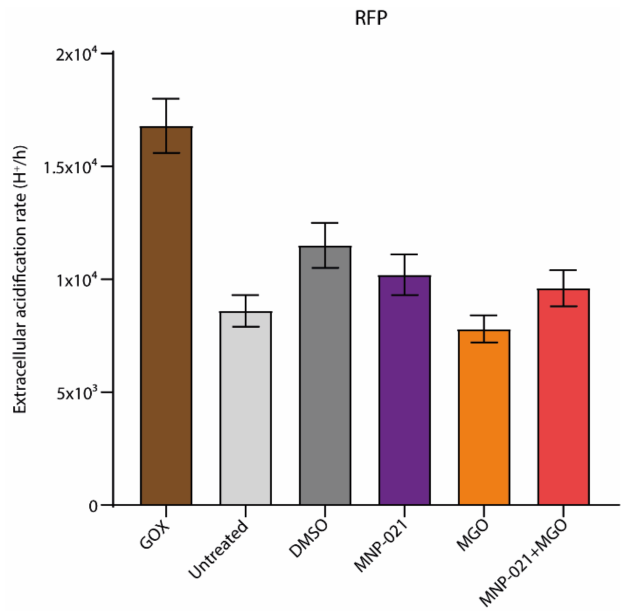


**Supplementary Figure 2.** GOX produced the highest acidification values, confirming the sensitivity of the assay.

**Supplementary Table 4.** Mean ± standard deviation of three replicates for free respiration buffer (FRB, baseline), oligomycin + glucose (glycolytic stimulation), and 2-deoxyglucose (DG) + glucose. Cells were treated with 1% DMSO, 10 µM MNP-021, 400 µM MGO, and the combination of MGO and MNP-021. GOX was included as a positive control to validate assay sensitivity, and untreated cells served as a reference for basal acidification.

| Treatment | FRB (Baseline) | Oligo + Glucose | DG + Glucose |
|---|---|---|---|
| **DMSO** | 11 500 ± 1,000 | 12 800 ± 900 | 11 700 ± 800 |
| **MGO** | 7 800 ± 600 | 11 900 ± 700 | 8 400 ± 700 |
| **MNP-021** | 10 200 ± 900 | 12 900 ± 1 000 | 11 500 ± 850 |
| **MNP-021+MG** | 9 600 ± 800 | 7 800 ± 1 200 | 7 600 ± 900 |
| **GOX** | 16 800 ± 1 200 | — | — |
| **Untreated** | 8 600 ± 700 | 9 200 ± 600 | 7 800 ± 650 |

#### *MNP-021 prevents microglia inflammatory response*

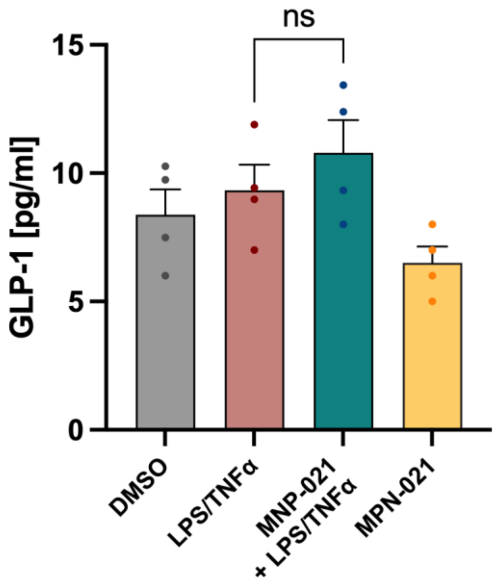


**Supplementary Figure 3. Effect of MNP-021 on GLP-1 release in HMC3 cells.** Microglia cells were pre-treated with MNP-021 (10 µM) and then stimulated with LPS/TNFα. No significant modulation of GLP-1 secretion was observed upon MNP-021 treatment. Data are expressed as mean ± SEM. ns, not significant.
