## Supplementary file 1 for "Unconventional Small Molecule MNP-021 Protects Neuronal and Glial Function from Diabetes-Associated Glucotoxicity and Neuroinflammation"

| Category | Term | Genes |
|---|---|---|
| GOTERM_BP_DIRECT | mitotic spindle organization | 9% |
| GOTERM_BP_DIRECT | mitotic cytokinesis | 9% |
| GOTERM_BP_DIRECT | positive regulation of mitotic sister chromatid separation | 6% |
| GOTERM_BP_DIRECT | neuron recognition | 6% |
| GOTERM_BP_DIRECT | positive regulation of mitotic cytokinesis | 6% |
| GOTERM_BP_DIRECT | positive regulation of attachment of mitotic spindle microtubules tc | 6% |
| GOTERM_BP_DIRECT | neuron migration | 9% |
| GOTERM_BP_DIRECT | mitotic spindle midzone assembly | 6% |
| GOTERM_BP_DIRECT | mitotic cell cycle | 9% |
| GOTERM_BP_DIRECT | positive regulation of mitotic cell cycle spindle assembly checkpoint | 6% |
| GOTERM_BP_DIRECT | cell division | 11% |
| GOTERM_BP_DIRECT | astrocyte differentiation | 6% |
| GOTERM_BP_DIRECT | cellular response to cytokine stimulus | 6% |
| GOTERM_BP_DIRECT | positive regulation of mitotic cell cycle | 6% |
| GOTERM_BP_DIRECT | cardiac muscle contraction | 6% |
| GOTERM_BP_DIRECT | regulation of signal transduction by p53 class mediator | 6% |
| GOTERM_BP_DIRECT | mitotic spindle assembly | 6% |
| GOTERM_BP_DIRECT | apoptotic process | 11% |
| GOTERM_BP_DIRECT | vasodilation | 6% |
| GOTERM_BP_DIRECT | G2/M transition of mitotic cell cycle | 6% |
| GOTERM_BP_DIRECT | roof of mouth development | 6% |
| GOTERM_BP_DIRECT | negative regulation of neuron projection development | 6% |

| Count | List Total | Pop Hits | Pop Total | P-Value | Benjamini | Fold Enrichment |
|---|---|---|---|---|---|---|
| 3 | 30 | 57 | 19478 | 0,00325 | 0,624 | 34,17 |
| 3 | 30 | 65 | 19478 | 0,0042 | 0,624 | 29,97 |
| 2 | 30 | 4 | 19478 | 0,00594 | 0,624 | 324,63 |
| 2 | 30 | 5 | 19478 | 0,00742 | 0,624 | 259,71 |
| 2 | 30 | 6 | 19478 | 0,0089 | 0,624 | 216,42 |
| 2 | 30 | 8 | 19478 | 0,0119 | 0,624 | 162,32 |
| 3 | 30 | 119 | 19478 | 0,0135 | 0,624 | 16,37 |
| 2 | 30 | 11 | 19478 | 0,0163 | 0,624 | 118,05 |
| 3 | 30 | 135 | 19478 | 0,0171 | 0,624 | 14,43 |
| 2 | 30 | 12 | 19478 | 0,0177 | 0,624 | 108,21 |
| 4 | 30 | 397 | 19478 | 0,0207 | 0,664 | 6,54 |
| 2 | 30 | 19 | 19478 | 0,0279 | 0,819 | 68,34 |
| 2 | 30 | 33 | 19478 | 0,048 | 1.00e+0 | 39,35 |
| 2 | 30 | 35 | 19478 | 0,0509 | 1.00e+0 | 37,10 |
| 2 | 30 | 43 | 19478 | 0,0621 | 1.00e+0 | 30,20 |
| 2 | 30 | 43 | 19478 | 0,0621 | 1.00e+0 | 30,20 |
| 2 | 30 | 47 | 19478 | 0,0677 | 1.00e+0 | 27,63 |
| 4 | 30 | 656 | 19478 | 0,0727 | 1.00e+0 | 3,96 |
| 2 | 30 | 51 | 19478 | 0,0733 | 1.00e+0 | 25,46 |
| 2 | 30 | 52 | 19478 | 0,0746 | 1.00e+0 | 24,97 |
| 2 | 30 | 67 | 19478 | 0,0952 | 1.00e+0 | 19,38 |
| 2 | 30 | 69 | 19478 | 0,0979 | 1.00e+0 | 18,82 |

| Bonferroni | FDR | Fisher Exact |
|---|---|---|
| 0,682 | 0,624 | 0,0000912 |
| 0,773 | 0,624 | 0,000135 |
| 0,877 | 0,624 | 0,0000137 |
| 0,927 | 0,624 | 0,0000229 |
| 0,957 | 0,624 | 0,0000343 |
| 0,985 | 0,624 | 0,0000638 |
| 0,992 | 0,624 | 0,0008 |
| 0,997 | 0,624 | 0,000125 |
| 0,998 | 0,624 | 0,00115 |
| 0,998 | 0,624 | 0,00015 |
| 0,999 | 0,664 | 0,00306 |
| 1.00e+0 | 0,819 | 0,000386 |
| 1.00e+0 | 1.00e+0 | 0,00118 |
| 1.00e+0 | 1.00e+0 | 0,00132 |
| 1.00e+0 | 1.00e+0 | 0,00199 |
| 1.00e+0 | 1.00e+0 | 0,00199 |
| 1.00e+0 | 1.00e+0 | 0,00237 |
| 1.00e+0 | 1.00e+0 | 0,0174 |
| 1.00e+0 | 1.00e+0 | 0,00279 |
| 1.00e+0 | 1.00e+0 | 0,0029 |
| 1.00e+0 | 1.00e+0 | 0,00476 |
| 1.00e+0 | 1.00e+0 | 0,00505 |
